# Identification of active modules in interaction networks using node2vec network embedding

**DOI:** 10.1101/2021.09.22.461345

**Authors:** Claude Pasquier, Vincent Guerlais, Denis Pallez, Raphaël Rapetti-Mauss, Olivier Soriani

## Abstract

The identification of condition-specific gene sets from transcriptomic experiments is important to reveal regulatory and signaling mechanisms associated with a given cellular response. Statistical approaches using only expression data allow the identification of genes whose expression is most altered between different conditions. However, a phenotype is rarely a direct consequence of the activity of a single gene, but rather reflects the interplay of several genes to carry out certain molecular processes. Many methods have been proposed to analyze the activity of genes in light of our knowledge of their molecular interactions. However, existing methods have many limitations that make them of limited use to biologists: they detect modules that are too large, too small, or they require the users to specify a priori the size of the modules they are looking for.

We propose AMINE (Active Module Identification through Network Embedding), an efficient method for the identification of active modules. Experiments carried out on artificial data sets show that the results obtained are more reliable than many available methods. Moreover, the size of the modules to be identified is not a fixed parameter of the method and does not need to be specified; rather, it adjusts according to the size of the modules to be found. The applications carried out on real datasets show that the method enables to find important genes already highlighted by approaches solely based on gene variations, but also to identify new groups of genes of high interest. In addition, AMINE method can be used as a web service on your own data (http://amine.i3s.unice.fr).

## Introduction

Current high-throughput technologies are now capable of reliably quantifying, at the scale of an entire organism, the molecular changes that arise in response to diseases or environmental disturbances. In order to extract from this pool of data the genes most related to the process under study (and therefore of most interest to the biologist), statistical methods are generally used to associate genes with numerical values reflecting the extent of their variation. In most studies, the genes considered of most interest are the ones whose relative differences in expression, or fold changes, are the largest. Unfortunately, the raw fold change is unreliable because it does not take into account the uncertainty inherent in gene expression measurements. To overcome this uncertainty, existing methods calculate a p-value to reflect the statistical significance of the variation.

Selecting the genes of interest on the basis of fold changes, p-values or a combination of both makes it possible to compile a list of genes whose expression varies most significantly. However, this procedure fails to identify genes whose combined action is essential in the process under study but whose individual scores are too low.

Though, as pinpointed by Rapaport *et al*. (2007), “a small but coherent difference in the expression of all the genes in a pathway should be more significant than a larger difference occurring in unrelated genes”. Arising from this observation, many methods have been proposed to analyze gene activity in the light of our knowledge about their molecular interactions. These pertinent sub-network are named “context-dependent active subnetworks” (He *et al*., 2017), “functional module” (Beisser *et al*., 2010), “maximal scoring subgraph” (Dittrich *et al*., 2008) or “altered subnetworks” (Reyna *et al*., 2018). The underlying idea is to identify a pertinent module of genes by simultaneously taking into account two criteria: one based on a measurement of genes activity and the other one reflecting the proximity between the genes in the module. One of the challenges is to define an appropriate scoring strategy based on these two criteria.

Nguyen *et al*. (2019) classified main computational methods for solving the active subnetworks identification problem in six categories: (i) greedy algorithms, (ii) random walk algorithms, (iii) diffusion emulation models, (iv) evolutionary algorithms, (v) maximal clique^1^ identification and (vi) clustering based methods. First two methods are simple and rapid but are highly dependent to the starting point of the algorithm that does not guarantee to reach global optima. Conversely, methods (iii) and (iv) are able to find global optima (in accordance with the scoring system used) or an approximation of it at the prize of a computational burden. Methods (v) do not fully answer the initial issue as it is probably not true that each gene involved in a biological process interacts with all the others. Finally, methods (vi) offers the advantage of being based on existing clustering algorithms, but these rely on a distance between objects that must be determined. Moreover, most of the clustering algorithms require to determine a priori the number of clusters to build, which is challenging. The commonality between all these methods is that their effectiveness is very dependent of the network topology. Unfortunately, it is known that molecular interaction networks are noisy and incomplete (De Las Rivas and Fontanillo, 2010). In recent years, network embedding (Cui *et al*., 2018) has proven to be a powerful network analysis approach by generating a very informative and compact vector representation for each vertex *v* in the network. The approach was initially considered as part of dimensionality reduction techniques (reducing for example a |*v*| × |*v*| adjacency matrix into a |*v*| × *m* matrix where *m* ≪ |*v*|). This dimensionality reduction allows to reduce noise and map nodes in a vector space in which distances between nodes accurately reflect their proximity in the original network. One of the most representative technique for network embedding is Node2vec (Grover and Leskovec, 2016).

Recent advances on deep learning has led to a plethora of methods based on deep neural networks for learning graph representations, methods that are often inspired by the learning of word embedding (Mikolov *et al*., 2013). Works on word embedding can be seen as learning linear sequences (word sequences). It has been shown that the resulting compact vector representations are capable of capturing rich semantic information about natural language. Processing graph structures is much more complicated. A popular approach is to convert a complex graph structure with a rich topology into a set of linear structures and then use a word embedding method to calculate the vector representation of each node.

As mentioned above, the identification of active modules requires the simultaneous consideration of two criteria. In existing methods, measurements of gene activity and their network proximity are either combined to form a single metric or optimized simultaneously using multiobjective algorithms (Corrêa *et al*., 2019). When working on embedded networks, the proximity facet is embedded in the vector space. It is then possible to focus on the detection of subspaces containing genes that have a high activity. Following this line, we propose AMINE (Active Module Identification through Network Embedding), a new and efficient method for active module detection based on Node2vec (Grover and Leskovec, 2016). Our method uses a greedy approach to build increasingly large clusters of nodes based on the similarity of their encoding vectors and to evaluate them according to a metric taking into account the activity of the contained nodes.

On artificially generated datasets, the effectiveness of the method compares well with several recent algorithms. On real datasets, AMINE allows to complement the results obtained with classical approaches by identifying new groups of genes of great interest.

## Results

### Evaluation of AMINE on artificial data generated by Robinson *et al*. (2017)

Many studies dealing with the identification of active modules have tested their methods on datasets generated by themselves and which are, at times, difficult to reproduce. Robinson *et al*. (2017) are among the few to give access to all materials used to test the MRF method they proposed. These materials contain the graph itself, the p-values associated with the nodes and the modules to be identified. It gives us the opportunity to apply our method on exactly the same data.

The simulated experiment used to evaluate the MRF method (Robinson *et al*., 2017) consists of a set of 1000 scale free graphs, each containing 1000 vertices associated with values simulated from a standard uniform distribution. In this dataset, there are three modules to discover (the hit modules), each containing 10 vertices. To simulate the fact that the vertices belonging to these modules represent differentially expressed genes, and are therefore associated with low p-values, these vertices are assigned simulated values from a truncated Gaussian distribution with mean 0 and standard deviation equals to 10^−6^. Robinson et al. (2017) compared their MRF method to NePhe (Wang *et al*., 2009), Knode (Cornish and Markowetz, 2014) and BioNet (Beisser *et al*., 2010) and report that MRF gives the best performance in term of recall. Since it is known that there are exactly 30 true hits in the dataset, the authors rate the different methods by considering only the proportion of true hits in each hit list of size 30.

We ran AMINE on these data to detect the three most significant modules. Since our method does not consider the size of expected modules as an input parameter, it can identify modules of various sizes. When considering only the recall criterion (proportion of hit nodes identified) as Robinson *et al*. did, AMINE is disadvantaged when it identifies small modules and favored when it identifies large modules. In order to have a fair comparison between AMINE and MRF, we apply the following rule: if the number of genes belonging to the three identified modules is greater than 30, then only 30 genes randomly chosen are considered. In the case where the three identified modules contain less than 30 genes, the recall obtained by our method is underestimated. Despite this disadvantage, the predictions made by AMINE obtained a slightly better recall than the one obtained by MRF (Fig. 1A, last boxplot ; the detail of the evaluation performed by Robinson *et al*. can be found in Fig. 7 of their article (Robinson *et al*., 2017)). The recall without truncation (number of true positive divided by the number of hit nodes), the precision (number of true positive divided by the number of nodes identified) and the F1 score (harmonic mean of precision and recall) were also plotted on the same figure. The median value of the F1 score is 0.76 while the minimum and maximum values are 0.31 and 0.94 respectively. The total number of genes in the three identified modules range from 18 to 46 with a median of 28 (Fig. 1B). This means that AMINE is able to identify modules close to the ground truth (although slightly smaller) without the need to specify their size a priori.

**Fig. 1.**
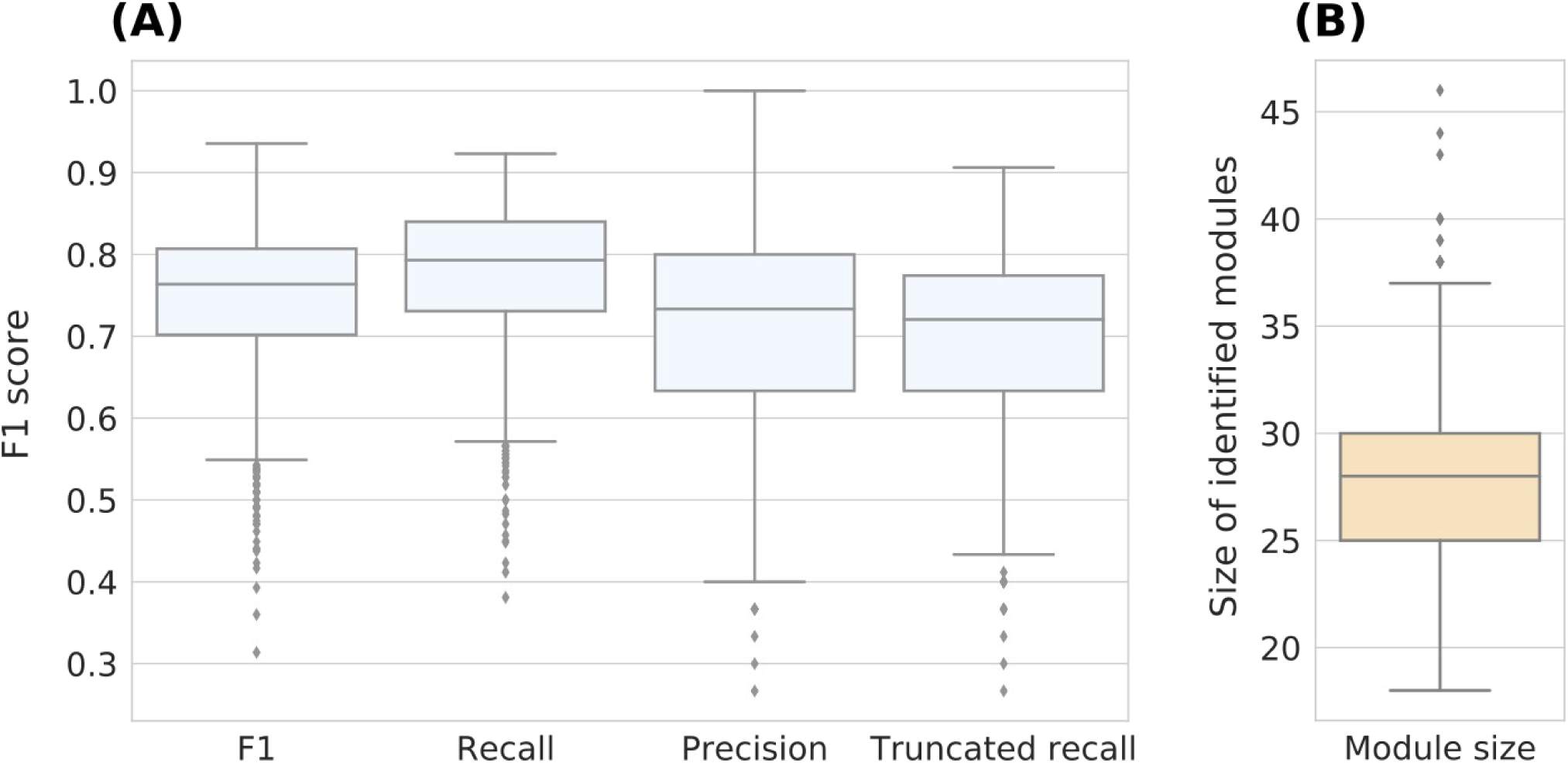
Evaluation of AMINE on artificial data generated by Robinson *et al*. (2017). **(A)** boxplots of the F1, recall and precision measures. The “trucated recall” displays the recall obtained by setting the maximum size of the 3 best identified modules to 30. **(B)** repartition of the size of the identified modules.

### Validation of the method on artificial dense networks

It has already been shown that simulating a biological network is a very difficult task (Pavlopoulos *et al*., 2011). However, we found that generating artificial graphs using Barabasi-Albert model of preferential attachment (Barabási and Albert, 1999), as performed in different studies (for example, the articles of Cornish and Markowetz (2014) and Robinson et al. (2017)) is too far from a real interaction graph for the results to be extrapolated (see supplementary Fig. S1 for an example of such sparse graph).

Using an extended version of the Barabási-Albert model of preferential attachment (Albert and Barabási, 2000), we generated several artificial networks with topologies relatively close to real interaction networks and one hit module to discover with size of 10 or 20 nodes (see materials and methods for details relative to the generation of dense networks and supplementary Fig. S2 for an example of a generated dense network). The performance of AMINE was compared with the methods COSINE (Ma *et al*., 2011), BioNet (Beisser *et al*., 2010), GiGA (Breitling *et al*., 2004) and a baseline consisting in simply picking the genes with the lowest p-values. The results are shown in Fig. 2 for networks comprising 1000 vertices and modules of size 10 and in Fig. 3 for networks with 1000 vertices and modules of size 20.

**Fig. 2.**
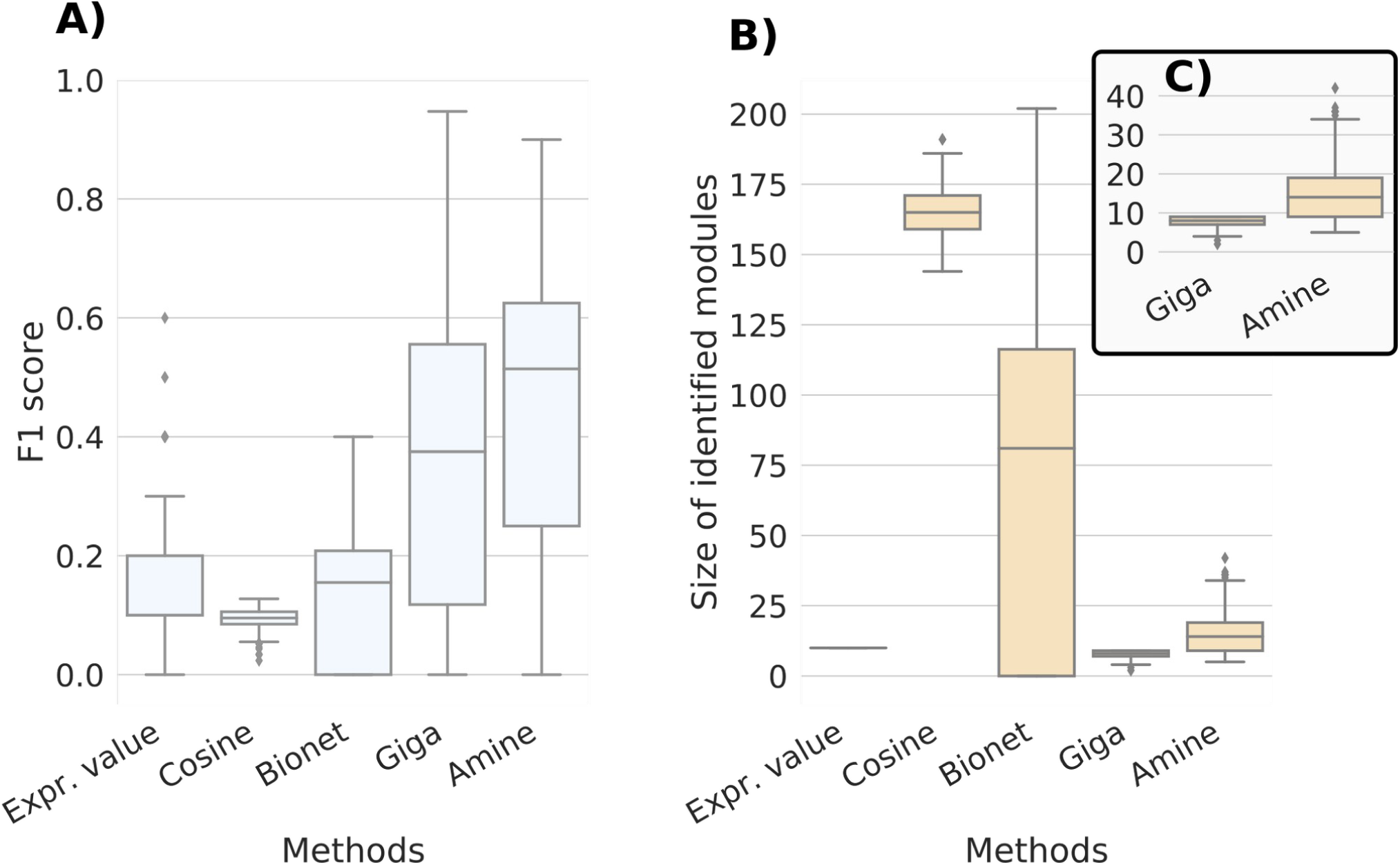
Results obtained for the identification of modules of size 10. Comparison of AMINE with COSINE, BioNet and GiGA on a task consisting in identifying a module of size 10 on a dense artificial network with 1000 vertices. The boxplots summarize the results obtained on 1000 different networks. “Expr. value”, that is used for baseline, consists in selecting the 10 genes with the lowest pvalues. **(A)** Boxplots representing the distribution of F1 scores. **(B)** Boxplots representing the distibution of identified module sizes. **(C)** Magnification of the distribution of module sizes for GiGA and AMINE.

**Fig. 3.**
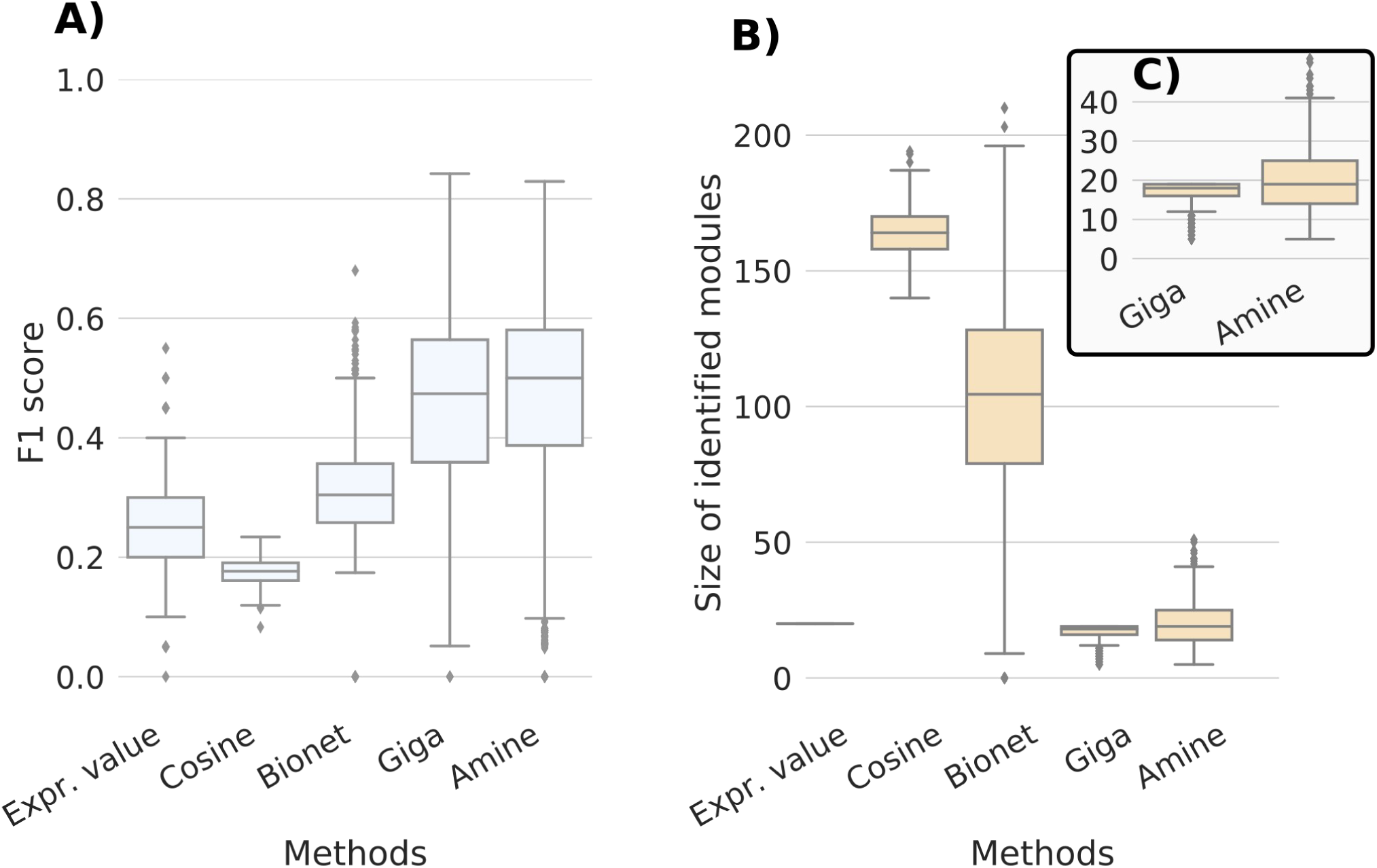
Results obtained for the identification of modules of size 20. Comparison of AMINE with COSINE, BioNet and GiGA on a task consisting in identifying a module of size 20 on a dense artificial network with 1000 vertices. The boxplots summarize the results obtained on 1000 different networks. “Expr. value”, that is used for baseline, consists in selecting the 20 genes with the lowest pvalues. **(A)** Boxplots representing the distribution of F1 scores. **(B)** Boxplots representing the distibution of identified module sizes. **(C)** Magnification of the distribution of module sizes for GiGA and AMINE.

We can observe that identifying module on denser network is a process much more complicated as the median F1 score drops from 0.76 on a sparse graph to values just above 0.5, depending of the size of the module. The scores of the other methods are much lower in the case of modules of size 10 ; some even score lower than the baseline. In the case of COSINE and BioNet, these poor results come with the prediction of large modules whose size exceeds 100 genes (Fig. 2B and Fig. 3B). For modules of size 20, the F1 scores of GiGA and AMINE are very close (Fig. 3B). However, it should be noted that GiGA uses a parameter that determines the maximum size of the module to be identified. In our experiment, we have set the maximum size to the size of the module to be identified, which, of course, helps the procedure. As shown in Fig. 2C and Fig. 3C, for GiGA, the size of the identified modules is always below the maximum. For AMINE, without any indication on the size of the modules being searched, we can see that the method predicts modules with a median size close to the ground truth size. This is a significant advantage of our method since biologists do not know in advance the size of the modules they are looking for.

### Scalability of the method

To test to what extent our method is able to scale to larger networks, we applied it to an artificial network of 10,000 vertices, that was generated using the same parameters previously defined. The processing time increases from 1 minute for a dense network of 1,000 nodes to 30 minutes for a dense network of 10,000 nodes. This processing time is acceptable as real biological networks are close to this size.

The distribution of the F1 scores obtained by AMINE as well as the distribution of the size of the identified modules are shown in supplementary Fig. S3-S4. For modules of size 10, the median F1 is in agreement with that obtained on networks of size 1,000. The overall performance is however less good because it is penalized by the fact that, in many cases, the module is completely missed (which explains why the second quartile starts at zero in the leftmost boxplot of supplementary Fig. S3). This can be explained by the fact that, as the number of nodes in the network increases, the number of non hit nodes that are associated with a random value higher than the values associated with hit genes increases. So, the probability that random modules score higher than the hit module increases too. This effect is less important for modules of size 20, for which the method works as well as for networks of 1,000 nodes. Regarding sizes, they are still close to the ground truth, although there is a greater spread of values for modules of size 20.

### Validation using a real gene expression dataset

In order to test the ability of AMINE to identify relevant biological functions, we downloaded from Gene Expression Omnibus a dataset relative to a study aimed at characterizing processes and genes associated to metastatic spreading in pancreatic ductal adenocarcinoma (PDAC). With a 5-year survival that has not significantly evolved for 30 years despite progresses in anticancer therapies (<6%), PDAC is a cancer with one of the bleakest prognoses of the most fatal cancers. In the study of Chiou *et al*. (2017), the authors compared two populations of primary PDAC cells according to the expression of HMGA2, a gene associated to a high metastatic potency and poor outcome in several cancers, including PDAC. Gene expression AMINE profiling was carried out from six pairs of HMGA2+/HMGA2-cell populations, each pair originating from PDAC primary tumors spontaneously generated in a genetically engineered PDAC mice model (PKC mice). The profiling generated 193 statically relevant modules (with associated p-values < 0.05) and we decided to focus on the five ones figuring at the top of the list to verify whether they were relevant to metastatic process in PDAC from an ontological point of view. The functions and signaling pathways associated to these five modules were explored using STRING database (Szklarczyk *et al*., 2019) and are shown in Table 1 and Fig. 4.

**Table 1.** Functions associated to the five best modules revealed by AMINE profiling of genes deregulated in PDAC metastatic HMGA2 positive cell population. Gene enrichment analyses were performed using data from the STRING database.

| Module number | P-value of the module | Function/Signaling pathway | Accession Id | P -value of annotation |
| --- | --- | --- | --- | --- |
| 1 | 2.24e-7 | Collagen fibril organisation<br>Extra cellular matrix organisation | GO:0030199<br>GO:0030198 | 4.48e-13<br>4.48e-13 |
| 2 | 7.47e-6 | HIF-1 signaling pathway<br>Central carbon metabolism in cancer | mmu04066<br>mmu05230 | 6.43e-10<br>1.17e-05 |
| 3 | 4.8e-5 | cell adhesion<br>ECM-receptor interaction<br>Focal adhesion<br>PI3K-Akt signaling pathway | GO:0007155<br>mmu04512<br>mmu04510<br>mmu04151 | 5.60e-06<br>1.54e-08<br>2.40e-07<br>1.36e-06 |
| 4 | 6.07e-5 | extracellular matrix structural constituent<br>Extracellular matrix organization | GO:0005201<br>MMU-1474244 | 1.82e-06<br>1.14e-06 |
| 5 | 6.17e-5 | transmembrane receptor protein tyrosine kinase signaling pathway<br>regulation of cell migration<br>regulation of peptidyl-tyrosine phosphorylation<br>angiogenesis | GO:0007169<br>GO:0030334<br>GO:0050730<br>GO:0001525 | 4.04e-16<br>4.64e-07<br>5.60e-07 |

**Fig. 4.**
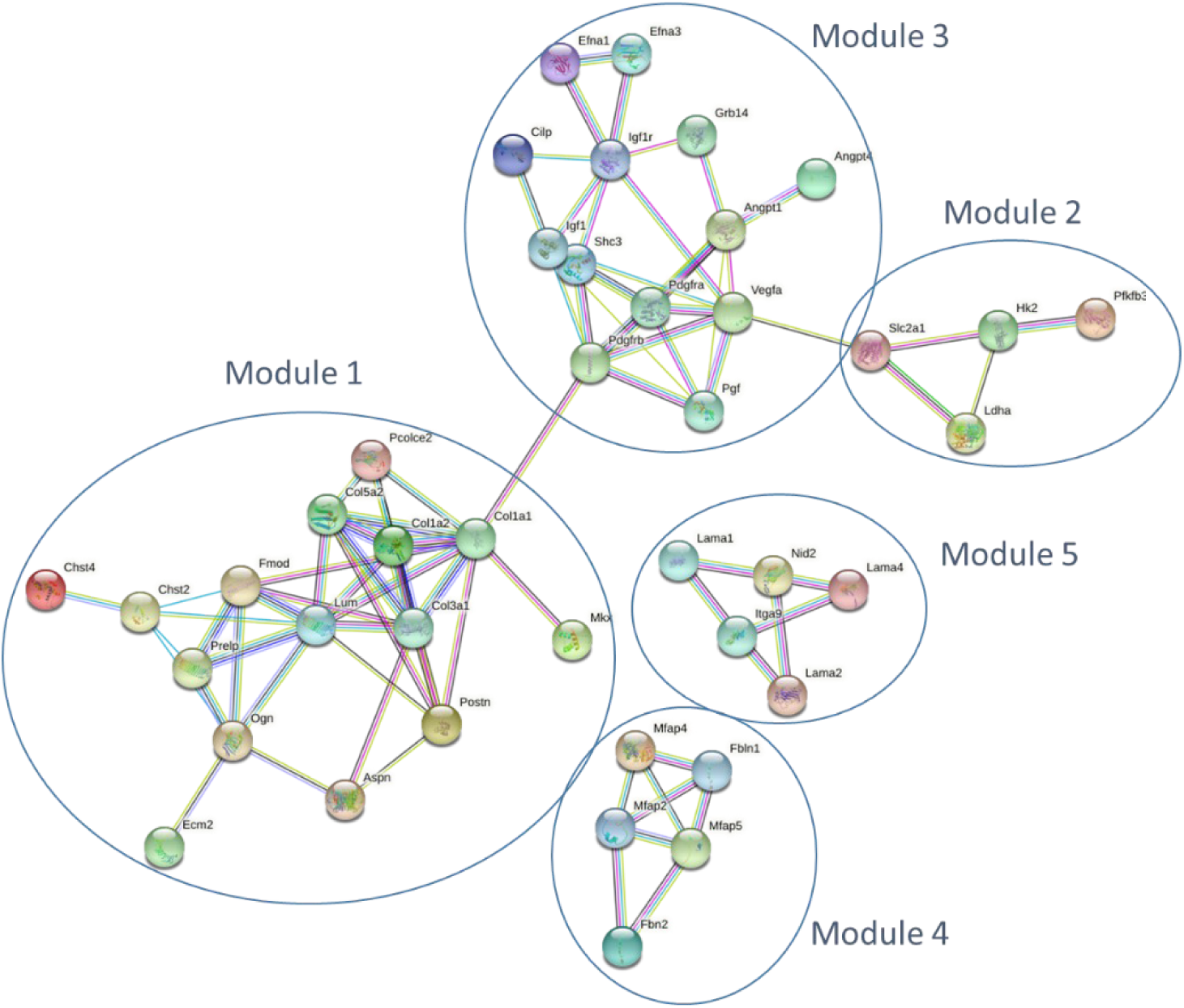
AMINE reveals modules associated to metastatic process in HMGA2 positive PDAC cells. The network was generated by STRING using the five first modules generated by AMINE from the list of deregulated genes in HMGA2+ PDAC cells available in Chiou *et al*., 2017.

### Extra cellular Matrix (ECM) organization and ECM cell interaction

A hallmark of PDAC is a pronounced collagen-rich fibrotic extracellular matrix (ECM) produced by fibroblasts and cancer cells, known as the desmoplastic reaction. The neoplastic epithelium exists within a dense stroma, which is recognized as a critical mediator of disease progression through direct effects of ECM on cancer cells (Hosein *et al*., 2020). Interestingly, three out of the five modules generated by AMINE were associated to ECM matrix organization (module 1 and 4) and cell interaction with ECM (module 3). Module 1 was more specifically linked to collagen fibril organization, one of the major constituent of PDAC ECM. Indeed, Collagen contributes to tumor cell aggressiveness, metastatic process and chemoresistance (Shields *et al*., 2012; Hessmann *et al*., 2020). Interestingly, Module 3 brings together genes involved in the regulation of cancer cell interaction with ECM through focal adhesion kinases and PI3K/AKT pathways (Fig. 4 and Table 1). Based on the available literature, these processes have been strongly involved in the aggressiveness of PDAC cell and the development of metastasis (Jiang *et al*., 2016; Jiang *et al*., 2020).

#### Response to Hypoxia

Extensive desmoplasia and hypo vascularization within PDAC results in significant intra-tumoral hypoxia (low oxygen) that contributes to its aggressiveness, therapeutic resistance, and high mortality (Hollinshead *et al*., 2020; Koong *et al*., 2000). Functional enrichment of modules 2 and 5 raised hypoxia-triggered functions, i.e. VEGF- and HIF-1-dependent pathways. These pathways drive angiogenesis, metabolism adaptation of cancer cell to hypoxia (Warburgh effect), cell cycle inhibition, enhanced migration and metastatic progression (Table 1 and Fig. 4). These results are in good agreement with the literature on PDAC; for example, these pathways are over-represented in genome-wide transcriptome profiling from ex-vivo human PDAC (Ghaderi *et al*., 2020; Shah *et al*., 2020). More interestingly, hypoxia-, VEGF- and HIF-1-associated pathways were repressed in PDAC cells in which BLIMP1, one of the most overexpressed gene in HMGA2+ cell subpopulation, was silenced (Chiou *et al*., 2017).

Altogether, these results validate our methods since non-oriented analysis of genes deregulated in pro-metastatic PDAC cells by AMINE retrieves gene modules involved in highly relevant functions in the context of the disease.

### In vitro functional validation of Blimp1-associated module in human PDAC cells

Blimp1 is one of the most overexpressed gene in pro-metastatic HMGA2+ PDAC cells (Chiou *et al*., 2017). In their study, Chiou *et al*. analyzed the consequences of Blimp1 silencing in mice PDAC cells to unveil its function in disease progression. Based on *in silico, in vitro* and *in vivo* experiments, the authors concluded that Blimp1 acts as a driver of the metastatic ability of PDAC cells. In particular, they found that Blimp1 is a Hypoxia/Hif-regulated gene in human and murine PDAC which is in a good agreement with functions recovered in modules 2 and 5 raised by AMINE processing (Table 1 and Fig. 4). Surprisingly, Blimp1 was not included in these modules, but was indeed detected in a module of 4 genes (module number 169 with a p-value of 0.044; Fig. 5A). Functional enrichment of this module unveiled functions associated to immune response, including regulation inflammation, interleukin production and Th17 cell differentiation (Table 2 and Fig. 5A). Interestingly, it is known that neoantigen expression in pancreatic ductal adenocarcinoma (PDAC) results in exacerbation of an inflammatory microenvironment that drives disease progression and metastasis (Hegde *et al*., 2020). It was therefore tempting to validate this result using a series of functional experiments. In this perspective, we first explored a RNA-Seq experiment performed by Chiou *et al*. revealing deregulated genes in Blimp1-silenced PDAC cells compared to control (Chiou *et al*., 2017). AMINE profiling of genes negatively regulated by Blimp1 silencing revealed 345 modules with associated p-values < 0.05. Among the ten best modules, we found that modules 2 (p-value < 1.01e-11) and 9 (p-value < 2.83e-8) were associated to cytokine production and inflammatory process (Table 3 and Fig. 5B) after functional enrichment. Next, in order to confirm the putative involvement of Blimp1 in epithelial cancer cell inflammatory process in vitro, we silenced BLIMP1 in MIA PaCa-2 cells, a human PDAC cell line, using siRNA silencing, (Fig. 6A), and explored how it modified the profile of cytokine secretion using a cytokine profiling array. Indeed, we found that Blimp1 repression triggered the production of IL-18Bpa and angiogenin, two anti-inflammatory factors (Lee *et al*., 2014) and reduced the secretion IL-6, a major pro-inflammatory interleukin (Tanaka *et al*., 2014) (Fig. 6B).

**Table 2.** Functions associated to the module embedding BLIMP1 revealed by AMINE profiling of genes deregulated in PDAC metastatic HMGA2 positive cell population. Gene enrichment analyses were performed using data from the STRING database.

| Module number | P-value of the module | Function/Signaling pathway | Accession Id | P -value of annotation |
| --- | --- | --- | --- | --- |
| 169 | 0.044 | regulation of interleukin-2 production<br>regulation of alpha-beta T cell differentiation<br>Th17 cell differentiation | GO:0032663<br>GO:0046637<br>mmu04659 | 0.0129<br>0.0129<br>0.00038 |

**Table 3.**
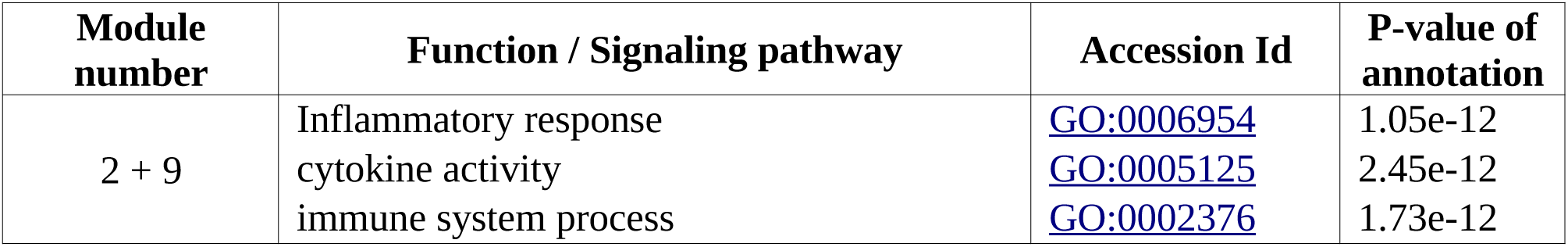
Functions associated to modules 2 and 9 resulting from AMINE profiling of genes down-regulated in PDAC cells silenced for Blimp1. The two modules were merged and subjected to gene enrichment analyses using data from the STRING database.

**Fig. 5.**
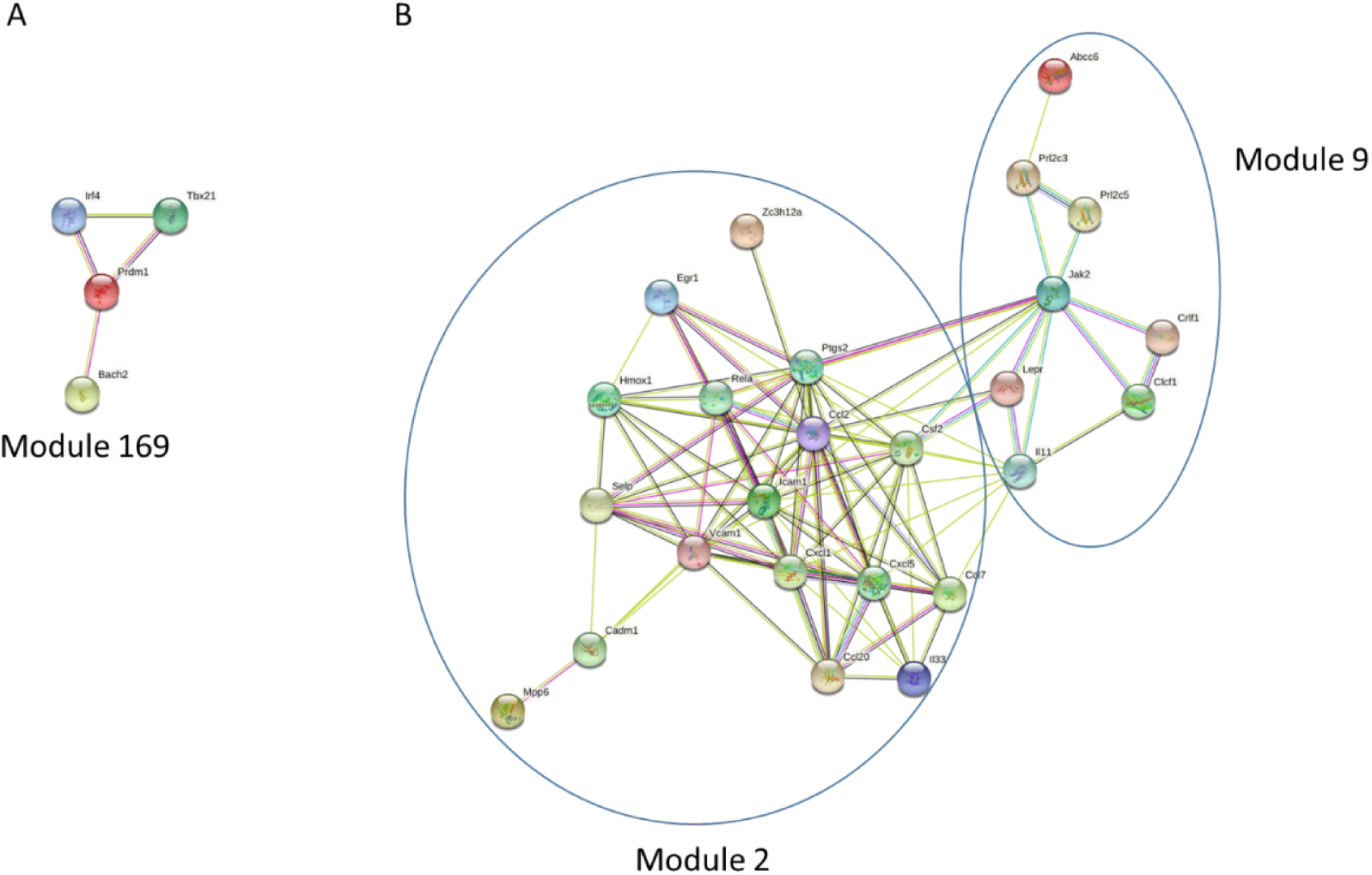
BLIMP1 is associated to immune response and inflammation in PDAC cells. **(A)** BLIMP1-associated module generated by the profiling of genes deregulated in PDAC metastatic HMGA2 positive cell population. **(B)** Modules 2 and 9 generated by the profiling of genes down-regulated in BLIMP1-silenced PDAC cells. The networks were generated by STRING.

**Fig. 6.**
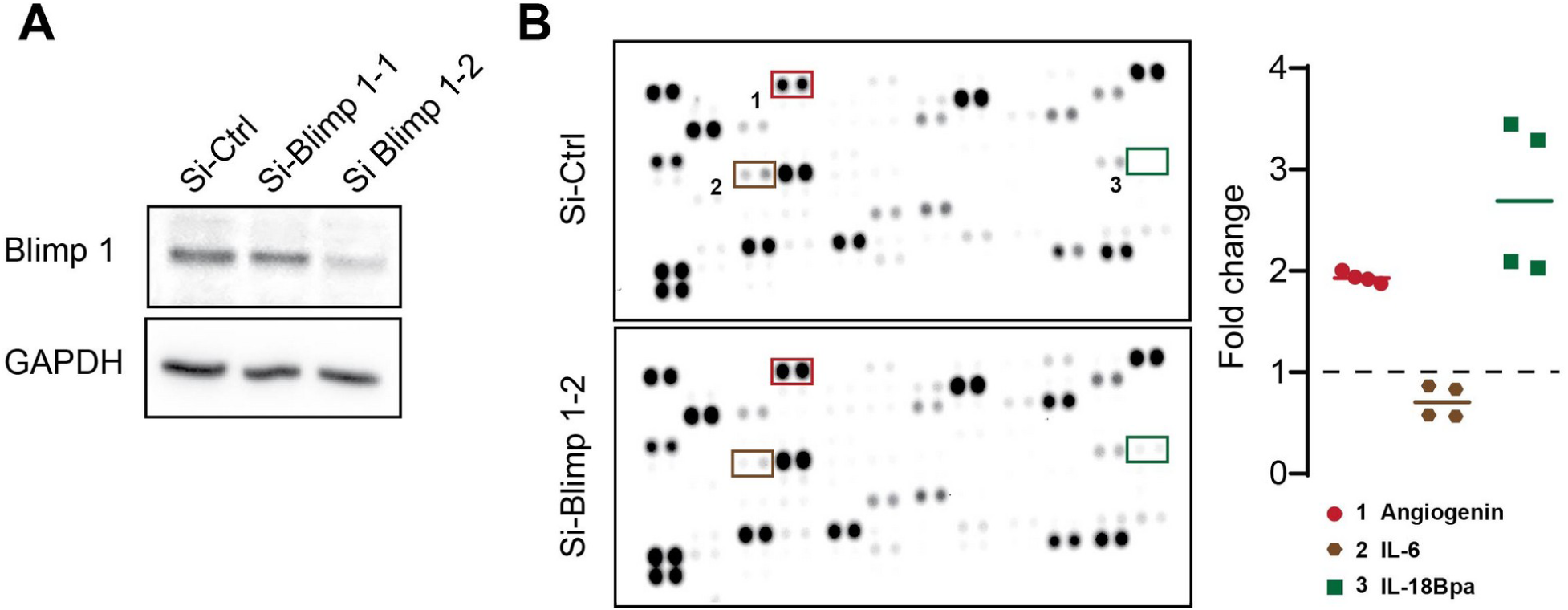
BLIMP1 silencing modify the cytokine secretion profile in PDAC cells. **(A)** Immunoblots of Blimp 1 in MIA PaCa-2 cells transfected with a non-targeting siRNA (Si-Ctrl), or with two different siRNA targeting Blimp-1 (Si-Blimp 1-1 and Si-Blimp 1-2). Data are representative of three independent experiments. **(B)** Soluble cytokine protein expression was assessed using cytokine arrays in si-Ctrl or si-Blimp 1 transfected MIA PaCa-2 (N=2; n = 4). Representative arrays are shown. On the left panel, values from densitometry quantification are shown as a fold change from the control.

Altogether, these results further validate AMINE as valuable method to detect relevant functional modules from large experimental datasets, and unveil a new function of Blimp1 in PDAC-related inflammatory process.

## Discussion

This paper proposes a new method for identifying gene modules that are activated as a result of a state shift caused by a biological experiment. Our method, called AMINE, uses as inputs, on the one hand, the ultimate result of any RNA-Seq analysis pipeline which is the differential expression of genes, and on the other hand, a network modeling the interactions between genes. Regarding the second point, the version of our method that is executable (at the address http://amine.i3s.unice.fr/) makes use of the protein interaction data provided by the STRING database for four model organisms: Caenorhabditis elegans, Drosophila melanogaster, Homo sapiens and Mus musculus. Thus, a user only needs to provide the differential gene expression data generated by the pipeline of his choice. From a very simple interface (supplementary Fig. S5), he only has to specify the name of the organism analyzed, the file on which the data are located and the Id of the columns containing the genes’ names, the p-values and optionally the fold changes to be able to launch the process. The address of the page containing the results is e-mailed to the user when the processing is completed. On the result page, the most significant modules are listed, however, all the modules found can be downloaded as an Excel document consisting of two sheets. The first sheet, named “list of modules” contains the list of all modules found. The results are presented in 4 columns containing the module number, the list of genes in the module, the s score of the module and the associated p-value. The second sheet, named “genes to modules”, is composed of two columns: the first one contains the name of a gene and the second one, the module to which it belongs.

Despite the large number of methods developed over the last 20 years, AMINE identifies, with a higher accuracy than its competitors, modules created computationally on datasets intended to mimic the topology of biological networks. Extrapolating these results to a measure of accuracy on real datasets is very difficult. There is no method to ensure that good predictions on artificial data translate into good predictions on real datasets. However, we have made a special effort to ensure that our simulations are close to real datasets. The networks we generate, with the parameters presented in this article, are closer to a real interaction network than the networks used by some competing methods.

However, these efforts run up against reality. Indeed, it is known that the interaction networks stored in public databases are both incomplete and contain erroneous interactions. In their study, Von Mering *et al*. (2002) estimate that, for Saccharomyces cerevisiae, the protein-protein interaction data (PPI) reported in public databases account for only one third of existing interactions. This observation leads us to believe that methods based very precisely on the topology of networks are not to their advantage. We think first of all of the methods based on the identification of cliques. Their performance is more than questionable if we consider that a large part of the interactions between proteins are unknown. If we consider that the missing interactions are randomly distributed on the graph, we can estimate that all the paths on the graph are impacted in the same way and thus that the methods based on random walks could be the least affected. Intuitively, we can indeed argue that, in a graph on which a certain proportion of the edges have been randomly deleted, if, from a source node A, random walks allow on average to reach node B before node C, then, on the complete graph, node B will probably always be closer to node A than node C. The other problem with methods based on graph traversals is that PPI networks are “small-world” networks, meaning that the neighbors of a given node are likely to be neighbors of each other, and most nodes can be reached from every other node by a small number of hops. Thus, any method that relies on graph traversals will find that a large portion of the network is close to any typical node (Cao *et al*., 2013).

Based on these considerations, we argue that network embedding methods can provide the backbone of a more reliable method by estimating distances between nodes that take into account the entire topology of the graph and, moreover, are little affected by the proportion of missing edges. Our method works on an embedding of an interaction network by adopting a greedy algorithm and an active subnetwork relevance measure defined in other papers (Ideker *et al*., 2002). The great advantage of our method is that it does not require any parameterization; it is not even necessary to indicate the number of modules to be identified or the size of the modules.

We have checked that our method performs well on artificial datasets and compares favorably with existing methods that are the current state of the art. We then processed a real dataset from a study focused on PDAC, on which AMINE retrieved modules associated with functions involved in PDAC metastatic process. Moreover, we highlighted functions that could not be detected using traditional approaches consisting in analyzing only the most differentially expressed genes. Our studies show that AMINE can identify modules corresponding to functions not revealed by traditional approaches. Indeed, we found that Blimp1, one of the most up-regulated gene in highly metastatic cells, participated in the regulation of pro-inflammatory process, a result confirmed by in vitro by the silencing of Blimp1 in human PDAC cells. However, we stress that our method is not an alternative to methods based on the identification of the most differentially expressed genes, but rather a complement to these approaches.

## Materials and Methods

The method AMINE predicts active modules from data consisting of a background knowledge about gene interactions and measurements representing, in the specific context of a given experiment, indicators of the involvement of genes in the studied process. This concept of gene-involvement is materialized by a p-value which quantifies, for each gene, the statistical significance of its variation (Fig. 7A).

**Fig. 7.**
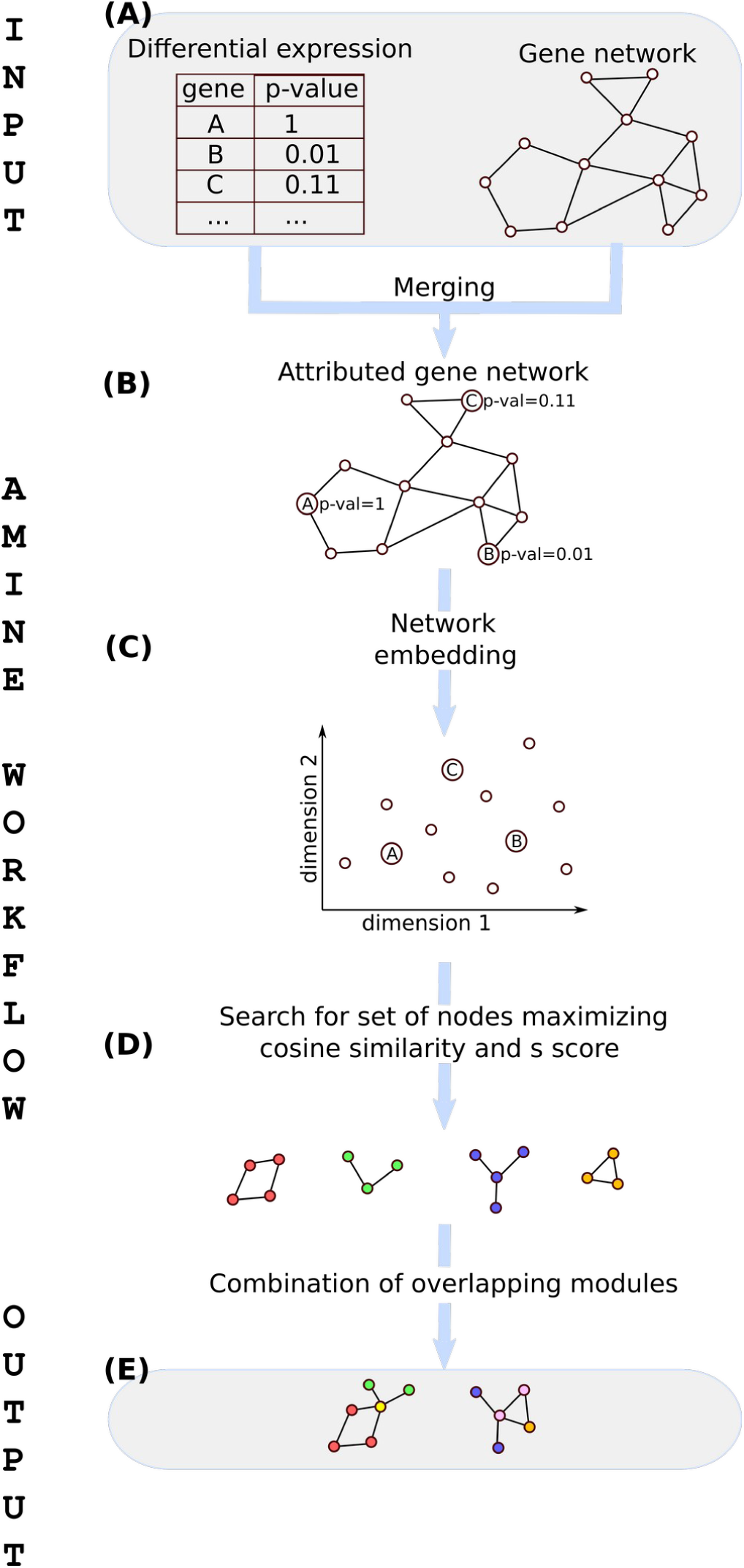
Workflow of the AMINE method. **(A)** Input data are composed of a table storing the significance of the expression variation of genes between two conditions and a network representing known gene interactions. **(B)** Data about gene interactions and gene variations are merged to generate an attributed gene network. **(C)** Nodes belonging to the attributed gene network are mapped to a low-dimensional space through the use of a biased Node2Vec method. **(D)** Sets of genes that are both cohesive and differentially expressed are identified in the embedded space by maximizing both the *s* scores of the nodes and the cosine distance cos between the vectors representing the nodes. **(E)** Redundancy in the content of modules is ruled out by combining sets of nodes obtained in the previous step while ensuring that the result remains spatially cohesive.

Data about gene interactions and gene variations are merged to generate an attributed gene network in which genes are annotated with a numeric attribute representing the extent of their variation (Fig. 7B). Mathematically, the dataset is represented as an attributed graph *G*=(*V, E, λ*) consisting of a set of vertices *V* (also called node, that symbolize the genes), a set of edges *E* ⊆ {(*u, v*) ∈*V*^2^ ∨ *u* ≠ *v*} and a value function *λ* (*v*): *V* → *R* which associates a value *p* ∈ *R* to each vertex *v* ∈ *V*. An induced subgraph of *G* is a subset *S* of the vertices of *G* together with those edges of *G* with both endpoints in *S*. Many active module detection algorithms focus on identifying induced subgraphs whose values associated with their nodes stand out from the values associated with the other nodes of the graph. We hypothesize that focusing heavily on the detection of connected sets of genes may not be optimal, given the fact that the interactions between genes described in the databases are still largely incomplete. For this, we adopt a definition of a module that is closer to the one used in cluster analysis: object that are grouped together (in a module) are more similar to each other than to those in other groups. The notion of similarity encompasses a component taking into account the distance on the graph between the vertices belonging to a module and a significant nearness between the values associated to these vertices.

### Scoring of a subgraph

Let *p*_*i*_= *λ*(*v*_*i*_) be the associated p-value of vertex *v*_*i*_. We aggregate the p-values associated to the nodes of a subgraph with Stouffer’s Z method; the same strategy used by Ideker *et al*. (2002). If we let *z* (*v*_*i*_)=*Φ*^−1^ (1 − *λ* (*v*_*i*_)), where *Φ* is the standard normal cumulative distribution function, then, the aggregate z-score *z*_*a*_ (*G′*) for an induced subgraph *G′* ⊆ *G* composed of *k* vertices, is computed with:

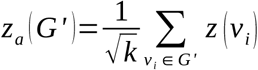

In order to get a subnetwork which has higher aggregation z-score compared with a random set of vertices, we define, still following the same methodology as Ideker *et al*. (2002), a corrected score *s* (*G′*) of a subgraph *G′* with:

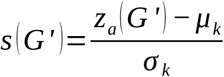

where the mean *μ*_*k*_ and standard deviation *σ*_*k*_ are computed based on a Monte-Carlo approach, taking 10,000 rounds of randomly sampling a connected subgraph of *k* vertices from *V*. From *s* (*G′*), we can easily compute the probability of observing in *G*, a subnetwork of the same size as *G′* with a corrected score at least as extreme as the one observed. This is given by the the one-sided p-value: *pvalue* (*G′*)=1 −*Φ* (*s* (*G′*)).

### Network embedding

A network embedding method is a function *ψ* : *V* → *R*^*m*^ that associate to each vertex *v* of the graph a vector *d* of size *m*. Node2vec (Grover and Leskovec, 2016) uses a biased random walk procedure which efficiently generates diverse neighborhoods of a given node. Node contexts are then processed with the word2vec method (Mikolov *et al*., 2013b). Node2Vec uses two parameters to control the walks. Intuitively, these parameters control how the walk explores and leaves the neighborhood of starting nodes. They allow a tuning between outward exploration and local walking.

However, in our case study, it may indeed be interesting, instead of using biases that only considered the topology of the network, to use the data associated to nodes, i.e. the value of *p*. The idea is to bias the walk so that, when the walker is located on a node, transitions to nodes with similar values of *p* (Fig. 7C) are favored. As *p* represents a p-value, the walker will be encouraged to favor visits of correlated and anti-correlated genes. We have conducted many experiments by replacing the parameters proposed by Mikolov *et al*. (2013) with our suggested use of similarity between nodes or by combining the different ways to bias the walk. It turns out, in the end, that using only the bias based on the similarity of p-values gives the best results. The bias we introduced allows to control the walk by assigning a transition *t* from a node *i* to a node *j* proportional to:

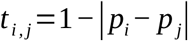

Other parameters tuning the Node2Vec method are summarized in Table 4.

**Table 4.**
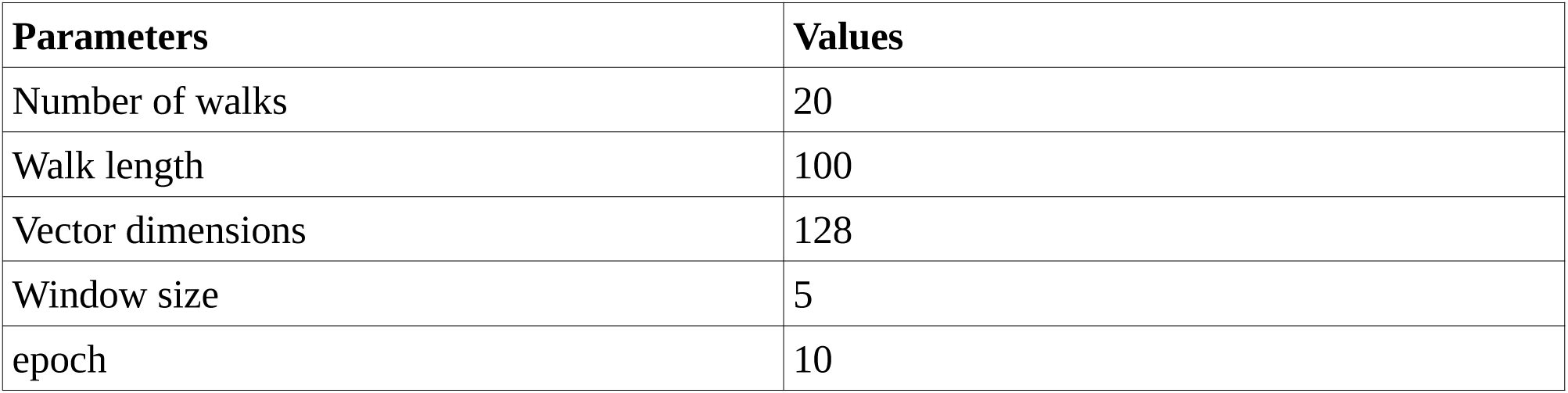
Parameters used for the Node2Vec method.

| Parameters | Values |
| --- | --- |
| Number of walks | 20 |
| Walk length | 100 |
| Vector dimensions | 128 |
| Window size | 5 |
| epoch | 10 |

### Algorithm

The cohesion measurement of a set of nodes on the graph can be determined on the embedded space using the cosine distance cos between the vectors representing the nodes. We use this property to identify, the most similar nodes to a given node *v*_*i*_ (represented as *similar* function in the algorithm summarized in Fig. 8). Thanks to a greedy approach, we collect, from each node, clusters *M*_*i*_ of increasing size evaluated using the *s* score previously defined. Our strategy is to expand the cluster as long as the *s* score increases (lines 8-10 of the algorithm). This allows to overcome the known drawback of the *s* score which is that high scores obtained by small sub-networks can be overtaken by random scores obtained in large networks (Nikolayeva et al., 2018). In practice, as we are very strict on the stopping condition, the clusters obtained are quite small (usually 5 nodes at most). At the end of this phase, we obtain a list of clusters, each one centered on a node, with each cluster being assigned a corresponding *s* score (Fig. 7D). The cluster centered on vertex *v*_*i*_ is thus denoted *M*_*i*_ with *M*_*i*_ ⊆ *V* and *v*_*i*_ ∈ *M*_*i*_.

**Fig. 8.**
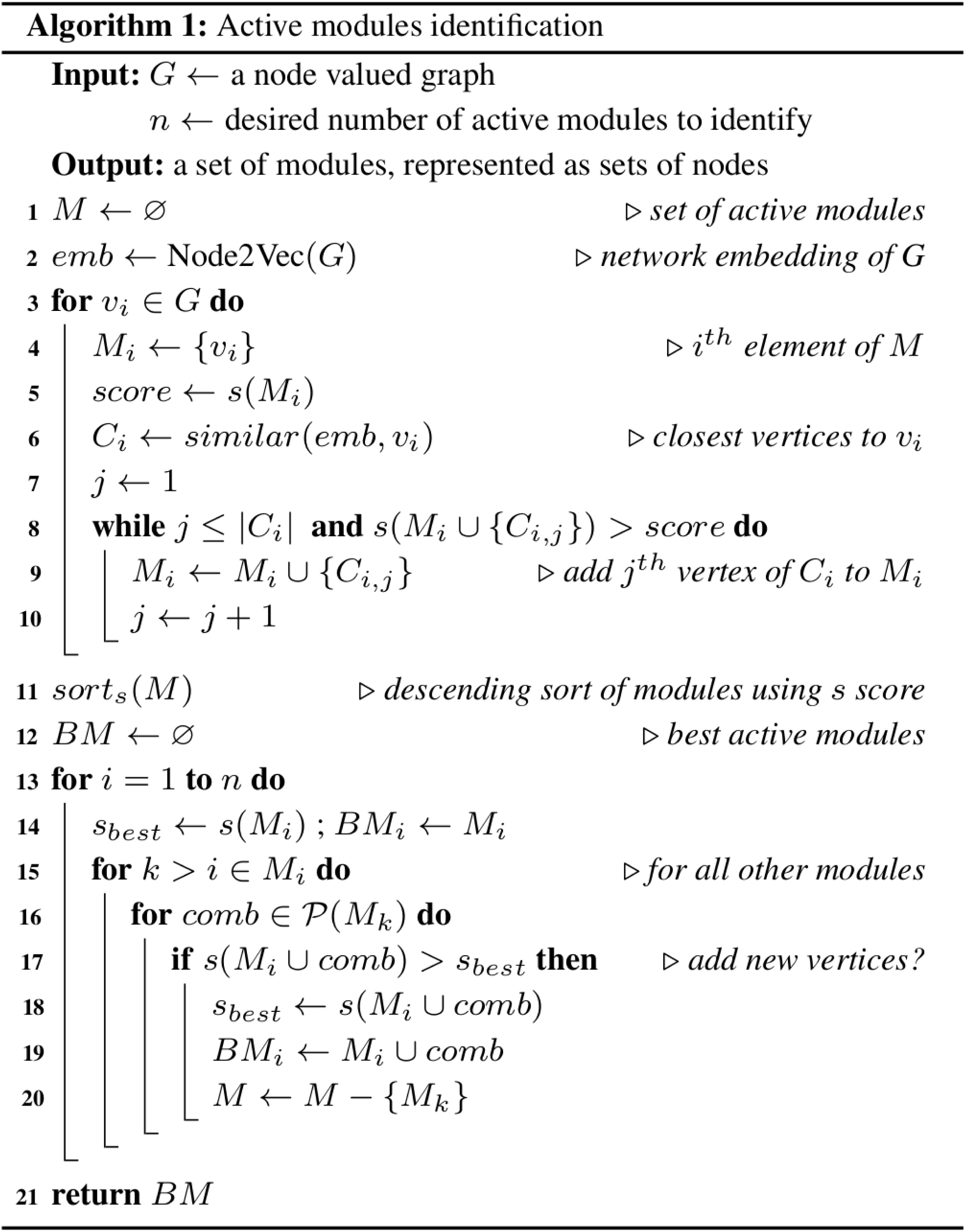
Algorithm of the method AMINE.

The next phase of the method consists in combining the different clusters while ensuring that new merged clusters remain spatially cohesive (Fig. 7E). We say that two clusters *M*_*i*_ and *M*_*j*_ are spatially cohesive when *M*_*i*_ ∩ *M*_*j*_ ≠ ∅. Starting from the module with the higher score (line 14 of the algorithm), the process consists in evaluating all possible clusters formed by *M*_*i*_ ∪ ℘ (*M*_*j*/*j* ≠ *i*_) using the *s* score, with ℘ (*M*_*i*_) denoting the powerset of *M*_*i*_ and keeping the modules with the highest *s* scores (lines 15-20). The workflow of the AMINE method is presented in Fig. 7 and Fig. 8.

### Generation of realistic interaction network

We use an extended version of the Barabasi-Albert model of preferential attachment (Albert and Barabási, 2000), to generate several artificial networks by varying the parameters *p* and *q* controlling the probabilities to add and remove edges respectively as well as the parameter specifying the number of initial nodes. Our results suggest that using 3 initial nodes with parameters *p* and *q* set to 0.09 and 0.70 respectively allows to generate random network with topologies relatively close to real interaction networks (Table 5).

**Table 5.** Comparison of artificial and real biological network. Comparison of several metrics associated, on the one hand, an artificial dense network generated with the extended Barabasi-Albert model using 3 initial nodes and setting parameters *p* and *q* to 0.09 and 0.7 respectively and, on the other hand, a subnetwork containing the same number of vertices extracted from the STRING database.

| Metric | Artificial dense network | Real network |
| --- | --- | --- |
| Number of vertices | 5980 | 5980 |
| Alpha coefficient | 1.908 | 1.602 |
| R square | 0.876 | 0.924 |
| Mean of neighbor connectivity | 42.207 | 47.720 |

1000 graphs were generated using these parameters (an example of this kind of graph is given in supplementary Fig. S2). The strategy to specify the value of nodes is exactly the same at the one of Robinson *et al*. (2017). In order to be able to do a comparison with other methods, we generated only one module of designated hits. As AMINE is dedicated to the identification of relatively small modules (in order to focus on really relevant genes that can be investigated by biologists), we have targeted our tests on the identification of small modules of size 10 and 20.

### Cell culture

The PDAC cell line, MIA PaCa-2, was obtained from Richard Tomasini CRCM, Marseille, France and culture in DMEM (Gibco™, Life Technologies Limited, Paiseley, UK) supplemented with 10% FBS, and penicillin/streptomycin. Cells were maintained at 37° C in a humidified atmosphere (5% CO_2_). Cells were tested routinely for *Mycoplasma* contamination.

### siRNA transfection

siRNAs (Sigma-Aldrich) were used for BLIMP 1 silencing. Non-targeting (si-Ctrl: SIC001) or BLIMP 1-targeting siRNAs (si-Blimp 1-1: 5’CUUGGAAGAUCUGACCCGA3’; si-Blimp 1-2: 5’CCUUUCAAAUGUCAGACUU3’ were transfected in MIA PaCa-2 cells using Lipofectamine^®^ RNAiMAX (Invitrogen, Life technologies Corp., Carlsbad, California, USA) following the manufacturer’s instructions. The final siRNA concentration was 30 nM. The medium was changed 8 h after transfection and the efficiency of the transfection was assessed by western blot after 72h.

### Western blotting

Cells were lysed in RIPA buffer supplemented with Complete Protease Inhibitor Cocktail and PhosSTOP Phosphatase Inhibitor Cocktail (Roche Diagnostics GmbH, Mannheim, Germany). Lysate were centrifuged at 12 000 rpm for 15 min at 4°C and then protein concentration was quantified using Bradford assay. Protein lysate were subject to SDS-PAGE and transferred onto a PVDF membrane. Membranes were blocked with 5% low fat milk in Tris-buffer saline –tween (TBS-T) for 1 hour. Membrane were incubated in Blimp 1 antibody (Cell Signaling, diluted at 1:1000) overnight. Membrane were washed in TBS-T followed by incubation with Horseradish peroxidase-conjugated secondary antibody for 1 hour at room temperature (Sigma-Aldrich). Signal was then visualized using ECL reagent (Immobilon^®^ Western, Millipore, Burlington, MA, USA) and chemoluminescence detection system (fusion FX7 Edge; Vilber, Marne-la-Vallée, France).

### Human Cytokine Array

For the cytokine assay, the Proteome Profiler™ Human XL Cytokine Array Kit (R&D Systems, Minneapolis, MN, USA) was used. The array was carried out using 500µl of cell supernatants obtained by incubating MIA PaCa-2 in DMEM 0% FBS, 48h after siRNA (si-Ctrl or si-Blimp 1-2) transfection, following the manufacturer’s instructions. For analysis of cytokine arrays, the intensity of each spot was measured using ImageJ software. Background was removed from all values, and they were normalized to the positive control spots.

## Acknowledgments

The authors are grateful to the OPAL infrastructure from Université Côte d’Azur and the Université Côte d’Azur’s Center for High-Performance Computing for providing resources and support.

## Funding

UCAJEDI Investments in the Future project managed by the National Research Agency (ANR) under reference number ANR-15-IDEX-01

French National Research Agency (ANR) through the LABEX SIGNALIFE program (reference # ANR-11-LABX-0028-01).

## Author contributions

Conceptualization: CP, OS

Methodology: CP, DP, RRM, OS

Software: CP, VG, DP

Validation: CP, DP, RRM, OS

Investigation: CP, DP, RRM, OS

Resources: RRM, OS

Data Curation: CP, VG

Visualization: CP, VG, DP, RRM, OS

Supervision: CP, OS

Writing—original draft: CP, DP, RRM, OS

Writing—review & editing: CP, VG, DP, RRM, OS

## Data and materials availability

All data are available in the main text or the supplementary materials.

## Supplementary Materials

Figs. S1 to S5.

## Footnotes

1 a clique is a subset of vertices of an undirected graph such that every two distinct vertices in the clique are connected (Seidman and Foster, 1978).

