## Supplementary materials for "Identification of active modules in interaction networks using node2vec network embedding"

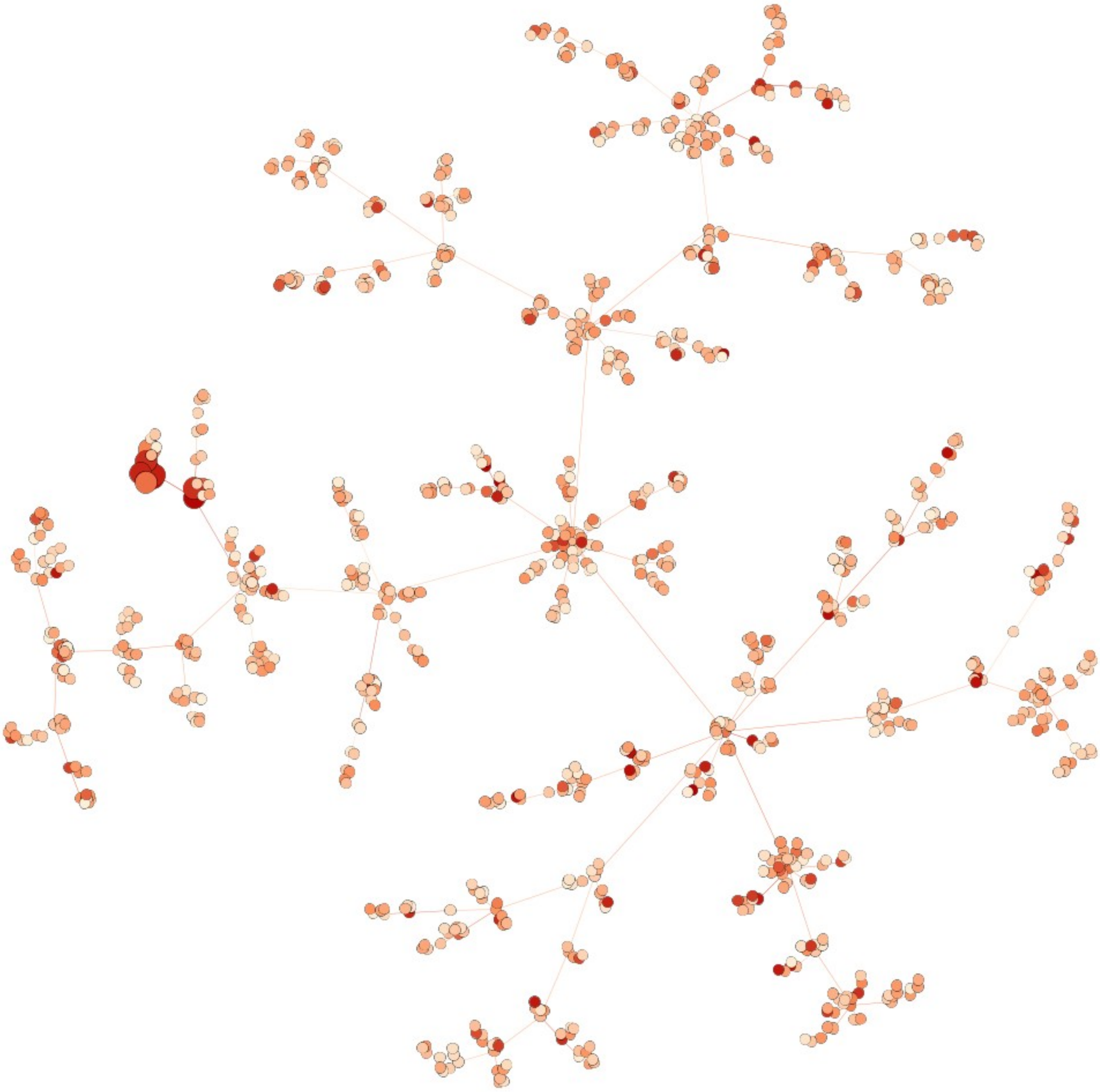

**Supplementary Fig. S1.** Example of a scale free graph of 1000 vertices generated with the Barabasi-Albert model of preferential attachment (Albert and Barabási, 2000). Node colors represents the values associated with nodes. Larger nodes constitutes the module to discover.

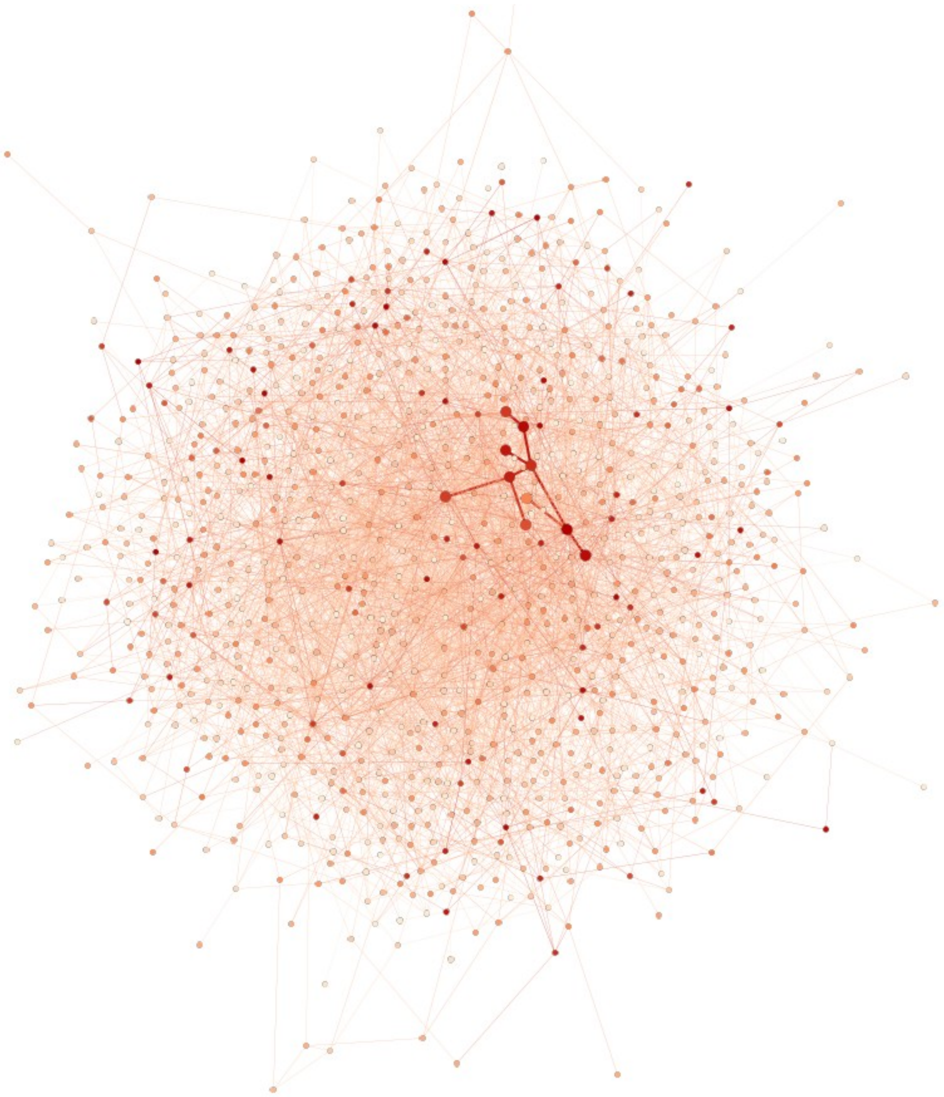

**Supplementary Fig. S2.** Example of a scale free graph of 1000 vertices generated with the extended version of Barabasi-Albert model of preferential attachment (Albert and Barabási, 2000). Node colors represents the values associated with nodes. Larger nodes, connected by wider edges, constitute the module to discover.

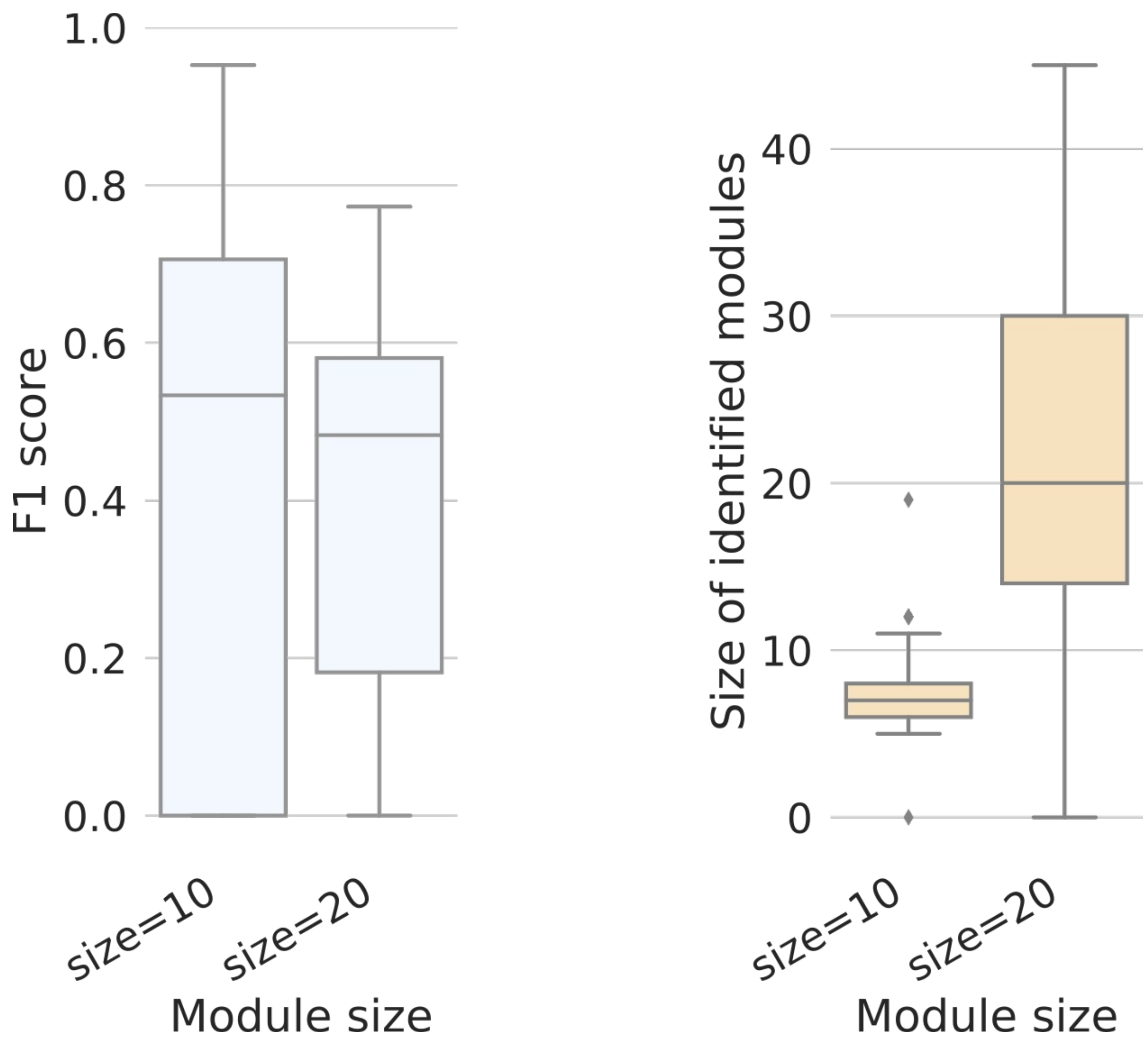


**Supplementary Fig. S3.** Boxplots representing the distribution of F1 scores and identified module size obtained by our method on dense artificial networks of 10000 vertices with modules of size 10 and 20 to discover.

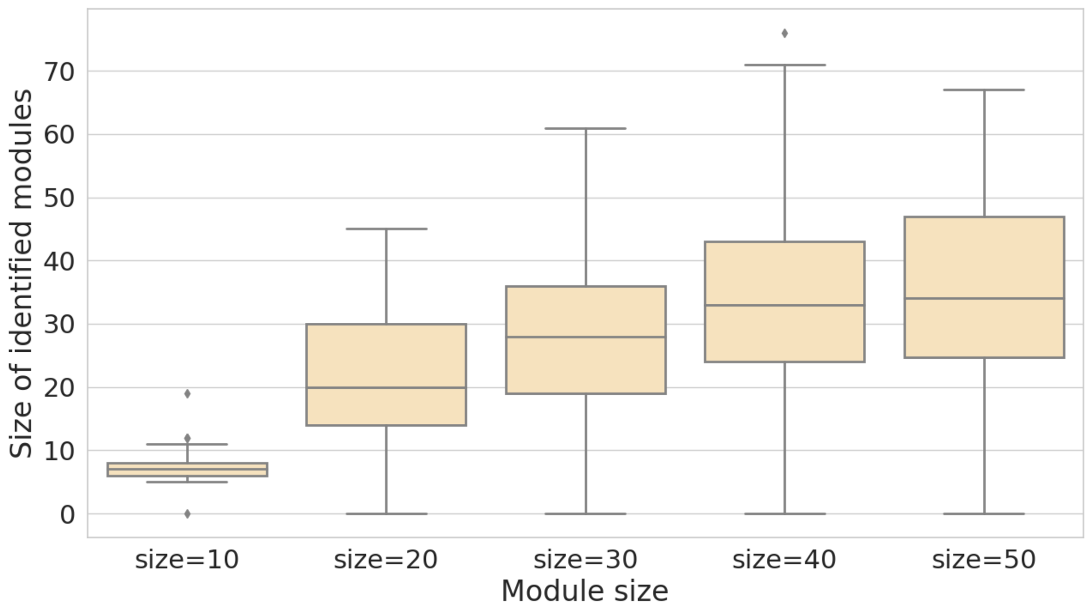


**Supplementary Fig. S4.** Boxplots representing the distribution of the size of the modules identified by our method on dense artificial networks of 10000 vertices for several theoretical module sizes.

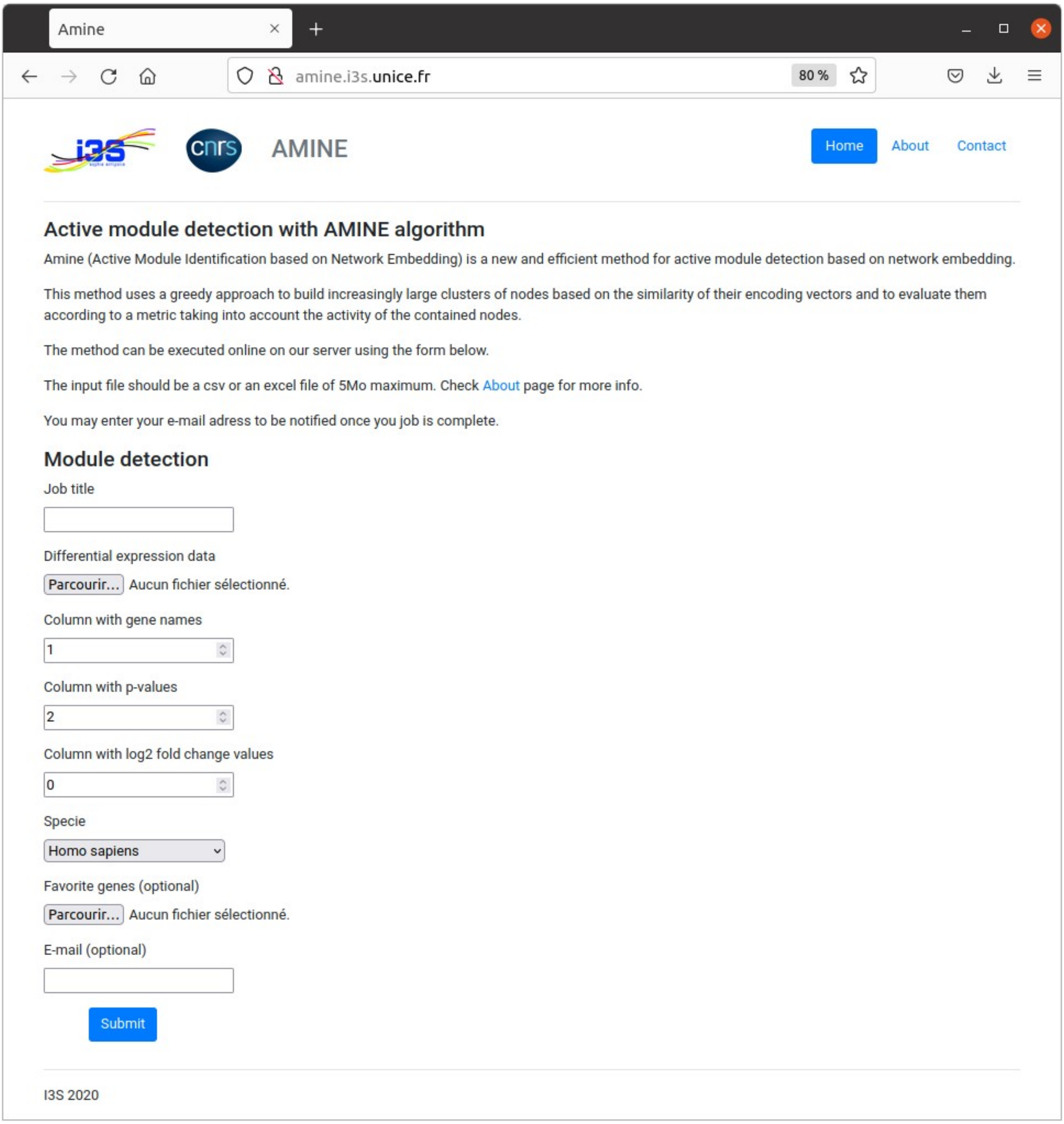


**Supplementary Fig. S5.** Input form of the AMINE method. P-values, representing the evidence of change in gene expression across experimental conditions, are specified in a file uploaded via the "Differential expression data" input field. At a minimum, the data consists of a tabular file with two columns: one containing the name of the genes and the other, the associated p-value. Another way to indicate the data to be used is to simply specify the result file produced by differential expression analysis methods such as DeSeq2 or edgeR. The input file can be in csv (comma-separated values) or xlsx format. The following two input fields are used to specify the columns containing the required data. The index of the column that contains the gene names must be specified in "Column with gene names" while that of the column with p-values must be specified in "Column with p-values". The index of the column with the log2 fold change can optionally be specified in the next field. Then, a drop-down list allows to choose the organism. It is possible to focus the search on modules containing specific genes. A file containing the list of these "favorite" genes (one gene name per line) can be specified with the "Favorite genes" input field. The e-mail address is used to notify the user of the completion of the task and to provide the address where the results are available.
